# The effect of trans-cinnamic acid isolated from the root cultures of the Baikal skullcap (*Scutellaria baicalensis*) on the life expectancy and survival of *Caenorhabditis elegans*

**DOI:** 10.1101/2022.04.12.488097

**Authors:** Lyubov Sergeevna Dyshlyuk, Margarita Yuryevna Drozdova, Vyacheslav Fedorovich Dolganyuk

## Abstract

Oxidative stress is an increase in reactive oxygen species, which are signaling molecules of various pathologies in the body of living beings. Baikal skullcap (*Scutellaria baicalensis*) is a valuable source of biologically active substances with various pharmacological activity. The aim of the study is to study the effect of trans-cinnamic acid isolated from root cultures of *Scutellaria baicalensis* on the lifespan of a model organism of *C. elegans*, as well as its survival under conditions of oxidative and thermal stress. It was shown that trans-cinnamic acid increased the lifespan of nematodes, while the best concentration of phytomaterials was 50 microns. In addition, all tested concentrations (10-200) had a positive effect on nematodes under oxidative stress caused by paraquat. No positive effect of trans-cinnamic acid was detected during thermal exposure. In general, the results show the antioxidant potential of trans-cinnamic acid from plant material under oxidative stress, as well as the positive effect of the substance on the lifespan of *C. elegans*.

## Introduction

Aging is a physiological, time-dependent process. The process itself is characterized by various damages that occur both at the cellular and molecular level. An example of disorders in living organisms is telomere depletion, mitochondrial dysfunction, genome instability. It is assumed that the number of people over the age of 60 will increase significantly and by 2050 will amount to about two billion [3]. This makes the problem of maintaining and strengthening the health of elderly people relevant, since aging is most often accompanied by various diseases, such as Alzheimer’s disease, Parkinson’s disease, cardiovascular diseases, cancer, diabetes.

Reactive oxygen species (ROS) are factors of damage to proteins, lipids, DNA and are highly reactive chemical forms that include oxygen. They are directly related to aging. The excess amount of these molecules in the body compared to antioxidants signals inflammatory processes, and also leads to cardiac dysfunction [12]. There is evidence of a negative effect of ROS in the development of neurodegenerative diseases [4].

Antioxidant therapy is a way of resisting oxidative stress. Natural antioxidants include biologically active substances (BAS) isolated from plant raw materials. They exhibit various properties: antioxidant, antibacterial, anti-inflammatory, anti-cancer, including geroprotective [1]. A plant such as Baikal skullcap (*Scutellaria baicalensis*) is an excellent source of secondary metabolites. This species is used in traditional Chinese medicine and grows on the territory of Russia, China, Korea, Japan. The plant includes about 126 low molecular weight compounds, 6 polysaccharides. At the same time, most BAS are obtained from aboveground parts or roots of the plant. The most famous flavonoids in the plant are – baikalin, baikalein, vogonin, vogonoside. The plant also contains essential oils and trace elements [1].

The *C. elegans* model organism is often used in various studies, including those related to aging. What makes this model popular is that it has a number of advantages: short life expectancy, small size, convenience in storage, anatomical features, the ability to study a number of diseases on it. Nematodes have various genes and pathways (the insulin/IGF pathway (IIS), the mTOR signaling pathway, dietary restriction) that modulate life expectancy and oxidative stress [5].

## Objects and methods

- Root cultures of the Baikal skullcap (Scutellaria baicalensis) were obtained at the Research Institute of Biotechnology, Kemerovo State University, Russia;
- Preparation of plant extract of the root cultures of the Baikal skullcap (Scutellaria baicalensis): Root cultures were crushed to a size of 1 mm. The sifted powder in an amount of 3 g was subjected to alcohol (70% ethanol) extraction at a temperature of 35 °C. To avoid evaporation of the solvent during extraction in a water bath, a reverse refrigerator was used. The total amount of extractant was taken 260 ml. Extraction was carried out twice. The holding time was 5 hours. The extract was analyzed by HPLC.
- Isolation of trans-cinnamic acid from the extract of the root culture of the Baikal skullcap (Scutellaria baicalensis): The preparative isolation of trans-cinnamic acid was carried out by concentrating the extract of the root culture of *Scutellaria baicalensis* at a temperature not exceeding 50 °C. The evaporated residue was dissolved in diethyl ether three times. After combining the ether fractions, the residue was evaporated on a rotary evaporator. Next, the evaporated mixture was dissolved in a solvent consisting of n-hexane and acetone. The fraction was chromatographed on silica gel with a mobile phase in the n-hexane-acetone gradient (1:0 to 0:1). After that, the flavonoid fraction was rechromatographed. The mobile phase consisted of n-hexane-chloroform (1:0 to 0:1). The eluate was dried. The isolated trans-cinnamic acid was controlled by the IR spectrum on the SF-2000 device (OKB SPECTRUM, Russia).
- Chemicals and reagents: The nematode growth medium (NGM) was prepared by adding 1 ml of 1 M sodium chloride, 1 ml of 1 M magnesium sulfate, 1 ml of 5 mg/ml cholesterol in alcohol, 25 ml of 1 M potassium phosphate to the sterile agar NGM. They were poured into Petri dishes of 20 ml each. The absence of bacterial contamination was confirmed by holding the cups for two to three days at room temperature. A solution of trans-cinnamic acid was prepared by adding dimethyl sulfoxide (DMSO). The concentration of the runoff solution was 10 M. The initial solution was diluted in sterile distilled water to obtain concentrations of trans-cinnamic acid of 100, 500, 1000, 2000 microns. 15 ml of solution was introduced into the hole with worms. As a result, concentrations of the substance 10, 50, 100, 200 microns were obtained. The solutions were stored at a temperature of 4 °C.
- Strain of Caenorhabditis elegans, content: The effect of isolated trans-cinnamic acid on nematodes was carried out using a wild strain of C. elegans. The N2 Bristol strain was grown on cups with NGM medium designed for growing nematodes. Previously, the NGM medium was seeded with an E coli OP50 culture in the form of a square, without touching the walls of the Petri dish. The cups were incubated for a day at 37 °C. Synchronous worms were obtained as described earlier [2]. Nematodes of stage L1 were cultured in liquid S-medium. Previously, a nocturnal culture of E. coli OP50 was added to the medium, the concentration of which was 0.5 mg / ml. Bacteria and nematodes were cultured in a 96-well plate in which 120 μl of suspension was kept for 48 hours at 20 °C. After that, 15 μl of 1.2 mM 5-fluoro-2-deoxyuredine (FUDR) was added to the wells and the tablets were left for 24 hours at 20 °C. After incubation, the worms were in the L4 stage.
- Life expectancy analysis: Assessment of the effect of different concentrations (0, 10, 50, 100, 200 microns) of trans-cinnamic acid was carried out in 96-well plates using a liquid S-medium. The experiment had a 6-fold repetition. Live and dead worms were counted every 4-7 days for 61 days. The study was completed when all the nematodes in the control died.
- Stress resistance analysis: The effect of trans-cinnamic acid treatment on the resistance of nematodes to oxidative stress was carried out in a liquid medium. 15 μl of trans-cinnamic acid was added to the worms in the tablets in various concentrations and the tablet was left to incubate at a temperature of 20 °C for 5 days. Next, live and dead nematodes were counted in each cell. After that, 15 μl of 1M paraquat was added to each cell of the tablet and incubation was continued in the thermostat at a temperature of 20 °C. Counting of live and dead nematodes was done after 24 hours and 48 hours. The results were compared between experimental and control groups of nematodes. The control groups were incubated without the addition of trans-cinnamic acid. The effect of trans-cinnamic acid treatment on the resistance of *C. elegans* nematodes to temperature stress was carried out in a liquid medium. After adding 15 ml of trans-cinnamon solution, the worms were incubated for 5 days at a temperature of 20 °C. Live and dead worms were counted in each cell and the tablet was transferred to incubation at 33 °C. After 24 hours and 48 hours, live and dead nematodes were counted. Control nematodes were incubated without the addition of trans-cinnamic acid.
- Statistical analysis: All experiments were performed in six independent trials. The analysis of statistical data was carried out using the Microsoft Office Excel 2007 software product. The statistical analysis of the obtained data was carried out using the Student’s simultaneous paired criterion, for each pair of interests. The differences were considered statistically significant at p < 0.05.

## Results and discussions

The results of identification of the obtained trans-cinnamic acid from the root cultures of the Baikal skullcap (*Scutellaria baicalensis*) are presented in Figure 1 in the form of an IR spectrum of an individual BAS with a degree of purification of at least 95%.

**Figure 1.**
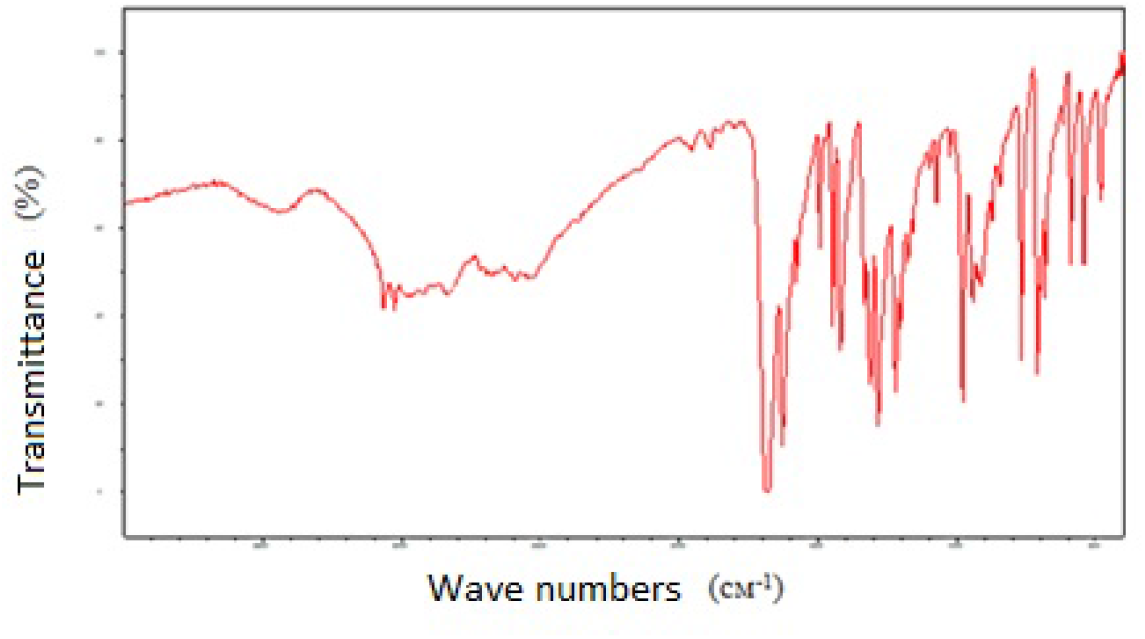
IR spectrum of trans-cinnamic acid

*C. elegans* is a popular model on which various drugs, including BAS, are tested [11]. By examining BAS, it is possible to determine both toxic doses of the substance and their concentrations, which positively affect the survival of worms. The effect of test doses of trans-cinnamic acid on average life expectancy is shown in Figure 2. It was found that after 8 days of incubation of nematodes with trans-cinnamic acid, all test concentrations positively affected the life expectancy of worms. The average life expectancy was increased by 18.1%, 26.3%, 24.1 by 36.6% at concentrations of trans-cinnamic acid of 50, 100, 200 microns, respectively. No positive effect compared to the control was recorded for the concentration of trans-cinnamic acid 200 microns from 13 to 34 days. This suggests that this dose of the substance shows little toxicity. From 34 to 61 days, better survival was observed for trans-cinnamic acid concentrations of 10, 50 and 100 microns compared to the control. A dose of 50 microns of trans-cinnamic acid showed a maximum survival rate of 9.8%.

**Figure 2.**
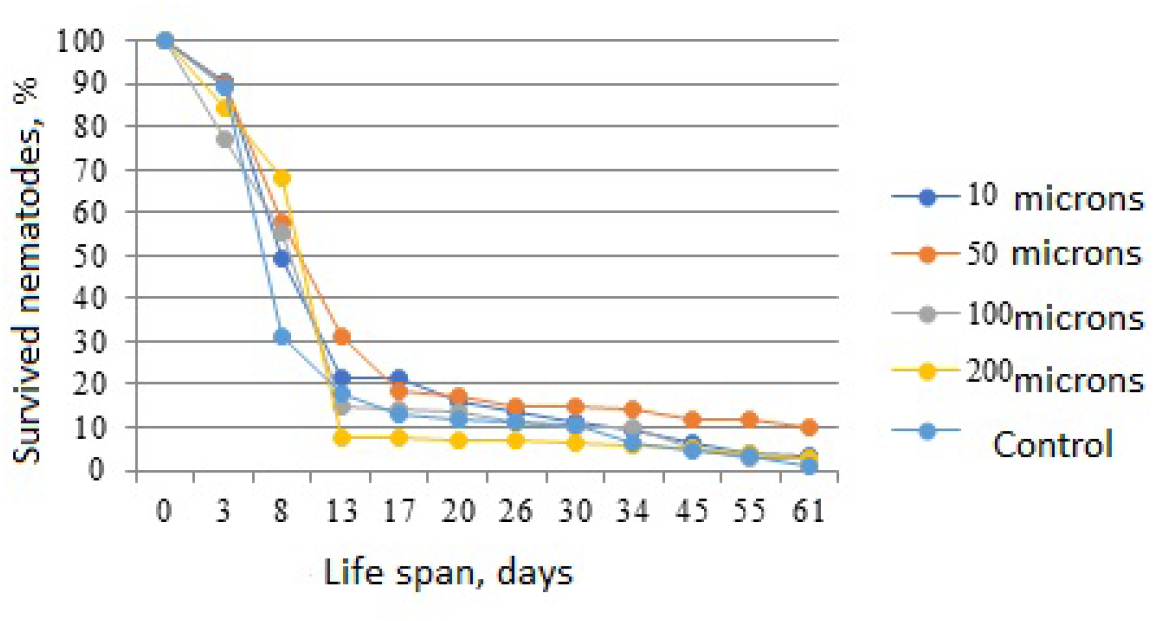
Effect of trans-cinnamic acid isolated from root cultures of Scutellaria baicalensis on the duration of wild-type worms

There are studies showing a positive correlation of life expectancy with survival under oxidative or thermal stress [6,9]. The effect of trans-cinnamic acid isolated from *Scutellaria baicalensis* root cultures on the survival of *C. elegans* under oxidative stress is shown in Figure 3. Paraquat as an inducer of intracellular ROS was used for analysis. The test substance had an effect on the stress resistance of nematodes to oxidative stress in the entire range of tested concentrations of 10-200 microns, compared with the control. The best result was observed when wild-type worms were treated with trans–cinnamic acid at a concentration of 100 microns - 94.3% and 81.8 after 24 and 48h incubation of nematodes with trans-cinnamic acid.

**Figure 3.**
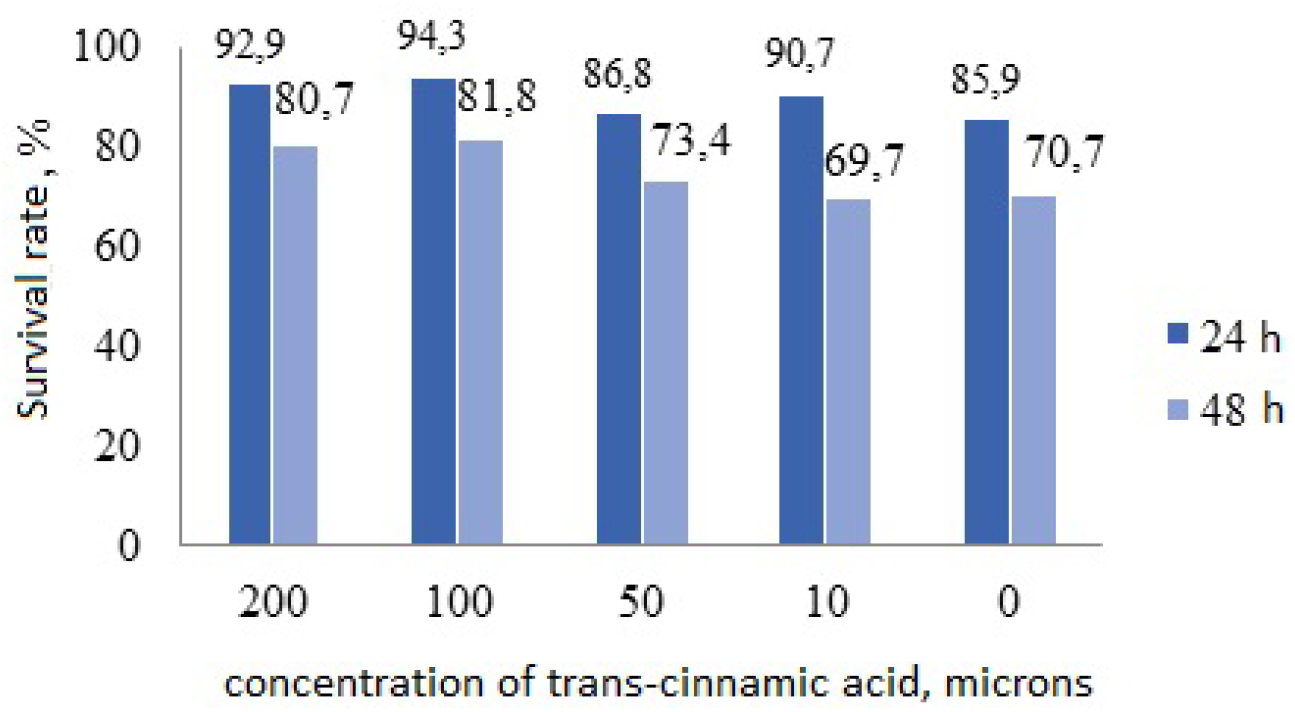
Effect of trans-cinnamic acid isolated from *Scutellaria baicalensis* root cultures on survival of wild-type worms under oxidative stress

Trans-cinnamic acid did not have a positive effect on *C. elegans* nematodes under heat stress. The highest concentration of trans-cinnamic acid 200 microns showed the worst result – 93.9 and 65.2% survival compared to the control – 96.3 and 80.6% (Figure 3).

**Figure 3.**
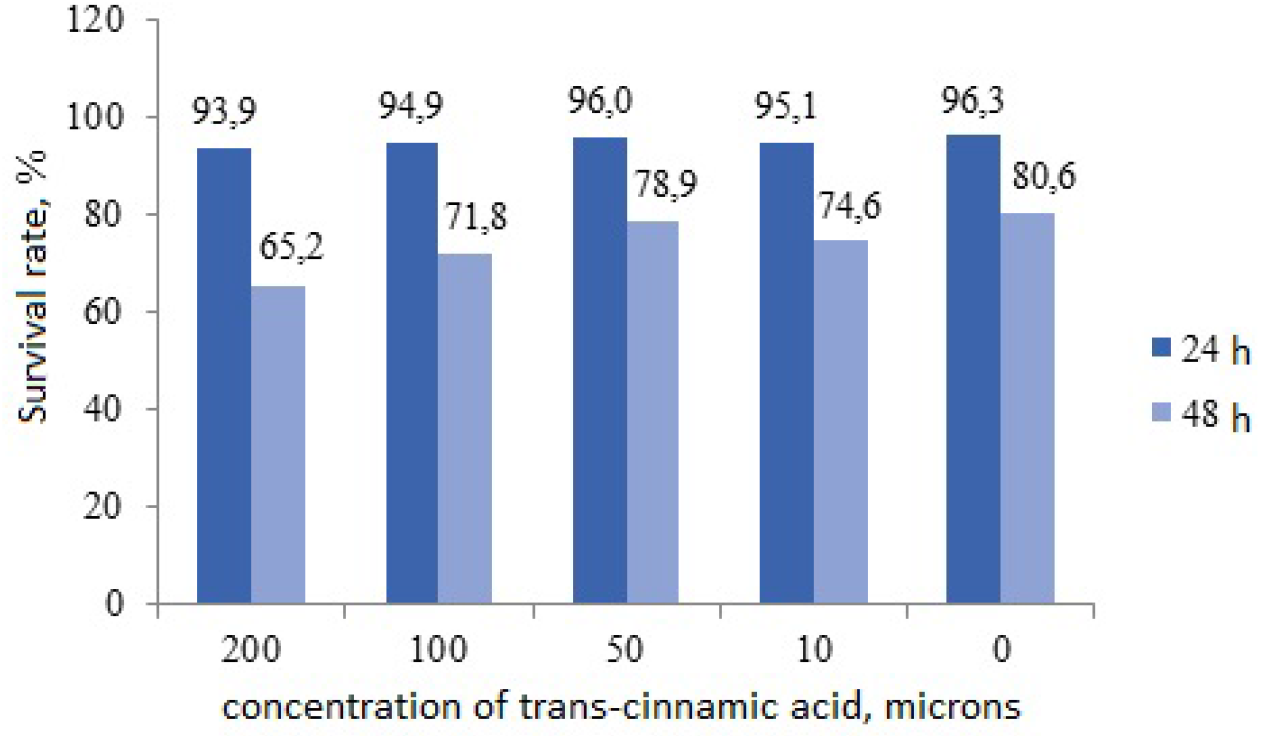
Effect of trans-cinnamic acid isolated from *Scutellaria baicalensis* root cultures on survival of wild-type worms under heat stress

Thus, studies show a positive effect of trans-cinnamic acid isolated from the root cultures of the Baikal skullcap (*Scutellaria baicalensis*) on the model organism of *C.elegans*. Trans-cinnamic acid increased the lifespan of nematodes, and also improves survival under oxidative, but not thermal stress. This shows that BAS can mitigate aging defects during oxidative processes in the organisms of living beings.

